# A morphing map model for place field organization in large environments

**DOI:** 10.1101/2024.04.19.590254

**Authors:** Michelangelo Naim, Mikail Khona, Christopher J. Cueva

## Abstract

Navigation and spatial memory, essential for animal survival, are founded on the hippocampal formation’s ability to host spatial neurons marking an animal’s position. Within controlled settings, the place cells of the hippocampus are activated in specific regions, the ‘place fields’. In this paper, we introduce a “morphing map algorithm” based on bat data from a 200-meter tunnel, highlighting a multifield, multiscale spatial representation of place fields. This algorithm posits that in vast environments, bats opt to select landmarks or ‘anchors’ for spatial representation. The method introduces a hyperbolic geometry in spatial representation, aligning with bat data from vast areas. We apply Procrustes distance to evaluate similarities between neural maps of bats and model-generated maps, revealing shared structure across bat maps, which might be dictated by common anchor points. Further explorations into anchor point ablation provide insights into their crucial role in map stability. The results present an evolved neural navigation strategy in expansive habitats and guide future spatial representation research across species.

## Introduction

Navigation and spatial memory are vital for the survival of animals in their natural habitats. The hippocampal formation houses various types of spatial neurons that represent an animal’s position and direction in space [1, 3, 4, 5, 6, 7, 8, 9, 10, 2]. Among these spatial cell types are “place cells”, hippocampal neurons that increase their spiking activity when an animal traverses a specific region of space, known as the neuron’s “place field” [1, 3, 11, 12, 13, 14, 15].

Individual place cells generally possess one or two place fields in smaller environments [3, 11, 16]. In contrast, multiple place fields are identified in upstream dentate gyrus neurons [16]. The majority of research on spatial representations in the mammalian brain has focused on rats and mice in small laboratory environments, typically using setups such as small boxes or short linear tracks around 1 to 2 meters in size. As a result, our understanding of spatial neurons in the hippocampal formation is primarily based on data from animals in confined laboratory settings.

Two studies have explored place cells on larger spatial scales [17, 18], but they used either a zig-zagging track of approximately 1-meter segments or a track passing through several small rooms. The largest single-compartment environment where place cells were recorded to date was less than 10 meters in size. However, outdoor navigation for mammals occurs in natural environments much larger than 10 meters. Wild rats, for instance, have been observed navigating more than 1 kilometer per night [19, 20]. Navigation across such distances demands spatial representation of vast environments, spanning hundreds of meters or kilometers [21].

Egyptian fruit bats fly nightly distances of up to approximately 30 kilometers to their preferred fruit trees, with fly-ways extending about 2 kilometers in width and 0.5 kilometers in height [21, 22]. Covering this space with typical place fields measured in the laboratory (around 10 to 20 centimeters in diameter, with one field per neuron) would necessitate around 10^13^ neurons, which is about 10^8^ times more neurons than the total number of cells in the entire dorsal hippocampal area CA1. Consequently, representing such large spatial scales with laboratory-sized place fields seems implausible. This discrepancy highlights a fundamental gap between laboratory-based navigation studies and natural, kilometer-scale outdoor navigation.

Recently, Eliav et al. [23] investigated bats flying in a 200-meter-long tunnel while recording the activity of hippocampal dorsal CA1 neurons using a custom wireless-electrophysiology system. They discovered that place cells recorded in the large environment displayed a multifield, multiscale representation of space: individual neurons exhibited multiple place fields with diverse sizes, ranging from less than 1 meter to over 30 meters, and the fields of the same neuron could differ by up to 20-fold in size.

### Morphing algorithm

How do bats organize their place fields in large environments? When a bat flies from position 1 to position *L*, it encounters multiple objects, or landmarks. We hypothesize that these landmarks serve as anchor points. The existence of place fields at other locations is determined by the activity of neurons at anchor points, following an algorithm based on morphing [24,25]. To model anchor points, we consider a set of binary patterns, *ξ*_*l*_, with the index *l* ∈ [1, *L*] representing the position of the anchor points in the trajectory.

One can construct place fields by initially selecting the activity of neurons at anchor point locations, with neuron sparsity matching experimental data. Specifically,

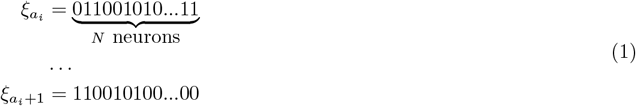

where 0 indicates the absence of a place field, and 1 denotes its presence.

To construct the sequence between all pairs of successive anchor points, we first choose the source 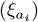 and target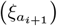 anchor points, where *a*_*i*_ and *a*_*i*+1_ represent the locations of neighboring anchor points. For approximately half of the neurons, the source and target states match each other 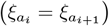 and the place fields of these neurons at the intervening locations are set to match the source and target values. If a neuron does not have the same values at the source and target anchor points, then we change the value from the source to the target value at a random intermediate location, ensuring that the target anchor point is obtained in the end.

Fig. 1c shows different maps with different anchor points. Interestingly, multiple maps have the same monotonic cumulative fraction of place fields (Fig. 1d). All the maps, with different anchor points, generate log-normal distribution of place field size and exponential distribution of the number of place fields (Fig. 1e).

**Figure 1:**
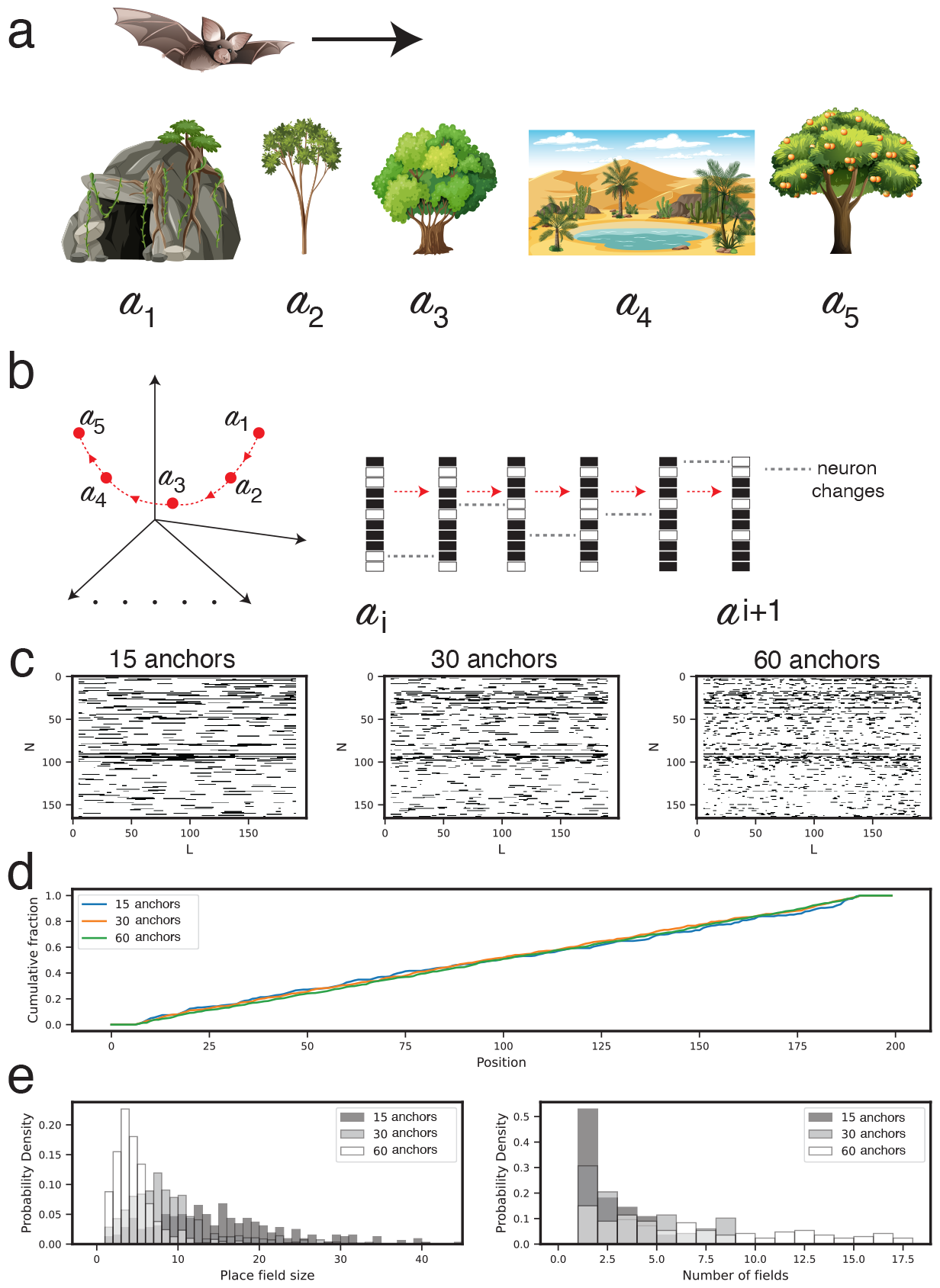
The morphing map algorithm for constructing place cell population codes for large 1D environments with anchors. (**a**) As the bat explores large environments, it chooses certain locations as “anchors” and stores them as specific patterns of place cell activity. (**b**) State space view of the population code (left) and visualization of N = 13 cells as the representation changes between 2 anchors *α*_*i*_ and *α*_*i*+1_. (**c**) Example morphing maps generated with varying number of anchors. The sparsity of each simulated neuron is matched to the sparsity of the data. (**d**) The centers of individual place fields remain uniformly distributed throughout the environment as the number of anchors varies. Hence the anchor locations cannot be read off from the place fields. (**e**) Variation of place field size (left) and number of fields per cell (right) as the number of anchor points in an environment vary.

In addition, we introduced the concept of measurement accuracy to account for the small discrepancy between our model and the observed data. The initial model showed a clustering of small fields around anchor points, a pattern not present in empirical data. This discrepancy was attributed to imperfect measurements. To rectify this, we introduced a 2% probability of field breakage, simulating the inaccuracies and resulting in a model that aligns more closely with the observed data (Fig. 2). The figure illustrates how varying levels of measurement precision impact the cumulative distribution of active place fields across different spatial maps. Specifically, Fig. 2a-b demonstrate the distinct influence of high (1.0) and slightly reduced (0.98) measurement accuracies on the distribution of both large and small place fields in relation to anchor points. Fig. 2c compares empirical data on bat place cell intervals with simulated distributions under these two measurement accuracies, thereby validating the hypothesis that slight inaccuracies in measurement can significantly alter the apparent spatial clustering of neural activity fields.

**Figure 2:**
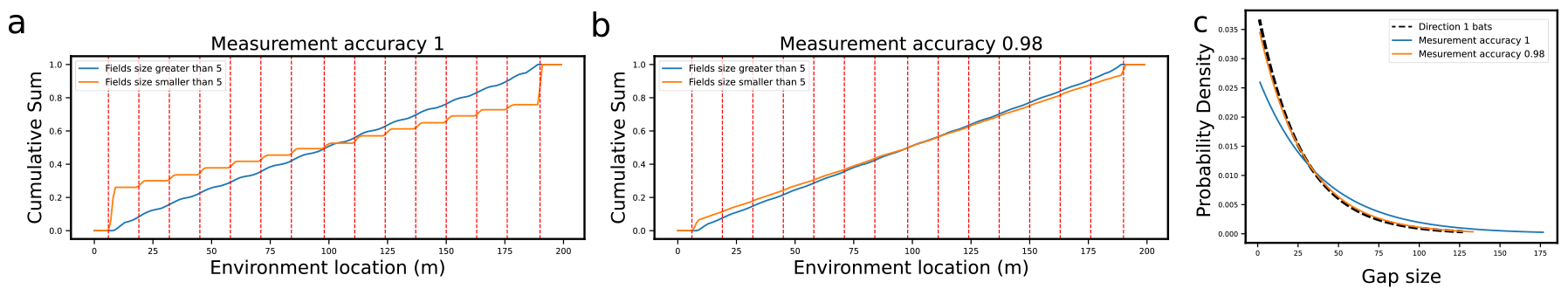
The influence of measurement accuracy. (**a - b**) **Impact of measurement accuracy on the cumulative sum of active fields for both big and small fields across 100 different maps with 15 anchor points**. Anchor points are denoted by dashed red lines. Here we present the hypothetical scenario when measurement accuracy is 1: the cumulative function for big fields (in blue) monotonically increases, showing no dependency on anchor points, while for small fields (in orange), it follows a step function near anchor points. (**b**) Here we show the situation when the measurement accuracy is adjusted to 0.98: the cumulative function of the small fields becomes smoother, closely resembling the curve for big fields. This alteration effectively simulates measurement inaccuracies and demonstrates the successful mitigation of small fields clustering exclusively around anchor points. (**c**) **Calibrating measurement accuracy using experimental data**. Here we show the distribution of interfield-intervals in the bat place cell data alongside hypothetical distributions under perfect measurement accuracy of 1 (in blue) and 0.98 (in orange). The observed distribution closely aligns with the scenario where the measurement accuracy is 0.98, validating our hypothesis that slight inaccuracies in measurement contribute to the small field breaks observed in the experimental data.

### Statistics of place fields generated by morphing map match those of bats flying in a long tunnel

To verify our hypothesis, we sought to compare the behavior of bats flying in large environments with the predictions of our morphing map model. The morphing map model reproduces the experimental observations quantitatively and qualitatively, which further supports our hypothesis that bats use a similar strategy for navigation.

In our model, the population codes are generated by morphing maps, induced by matching the sparsity of the neural activation patterns (Fig. 3a). The model predicts that the size of place fields and the number of place fields follow specific distributions: the size of place fields follows a log-normal distribution, and the number of place fields shows an exponential distribution (Fig. 3b).

**Figure 3:**
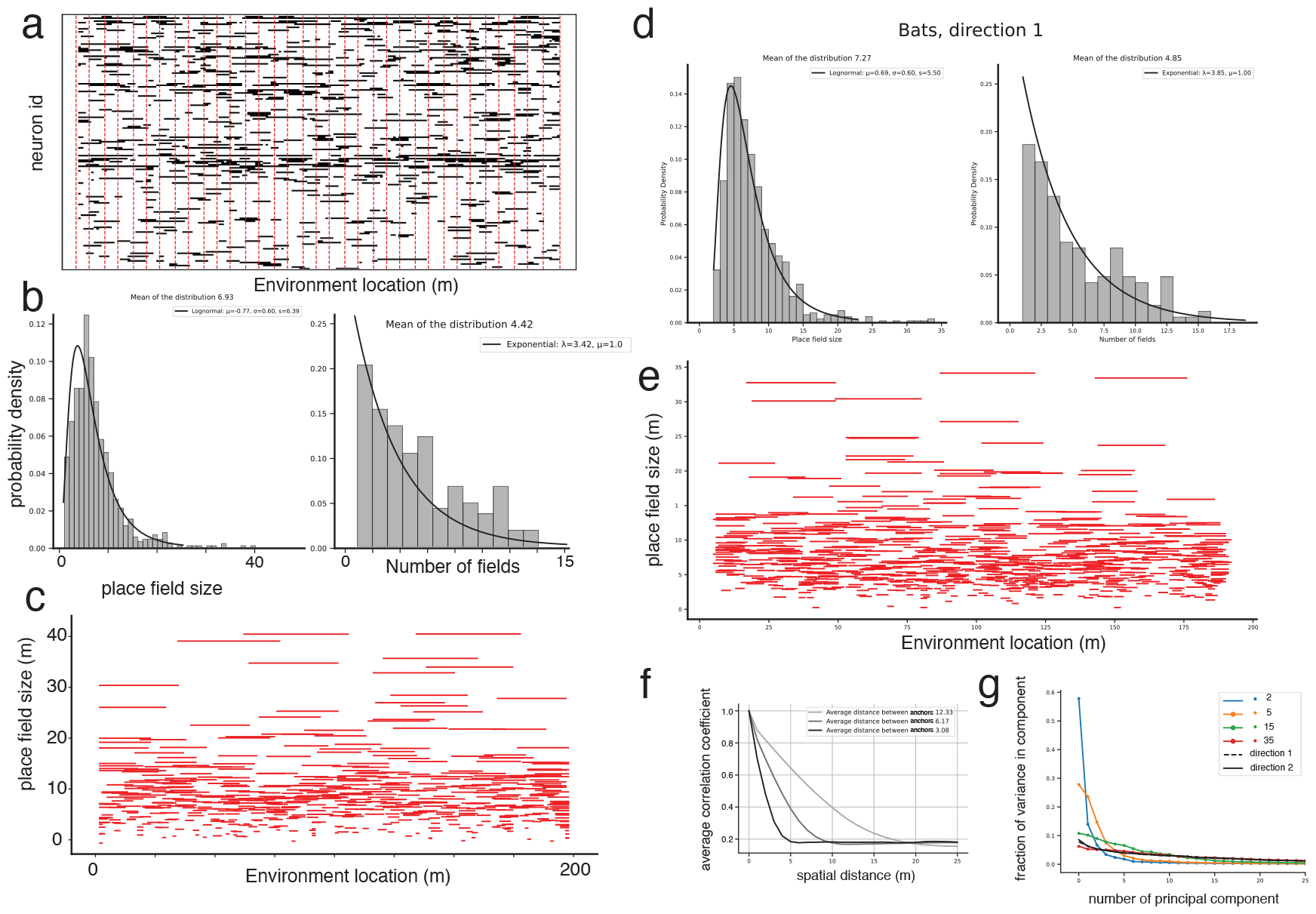
Population codes generated by morphing maps match quantitatively and qualitatively with bats flying in large environments. (**a**) Example morphing map population code induced by matching sparsity. The locations of anchor points are denoted by the red vertical lines. (**b**) Place field size and number of place fields from morphing map. (**c**) Ordering of individual place fields by their size shows a hierarchy, suggesting hyperbolic geometry. (**d**) Place field size and number of place fields for bat data. (**e**) Same as (c), but for bats. (**f**) Average spatial correlation for morphing maps with different anchors. (**g**) PCA of the population code matrix from morphing maps with 2, 5, 15, and 35 anchors and also for maps from bats showing that the geometry of the bat maps are consistent with high dimensional geometry of morphing maps with 35 anchors.

Moreover, when ordering the place fields according to their size, our model exhibits a clear hierarchy, which suggests a hyperbolic geometry underlying the representation of the spatial environment (Fig. 3c). This is a novel prediction of our model that requires experimental verification in future studies.

Remarkably, these distributions and the observed hierarchy in the place field size are also present in the data from bats flying in large environments (Fig. 3d-e). This demonstrates a close match between the predictions of our model and the experimental data, suggesting that bats might indeed use a morphing map mechanism to navigate large and gradually changing environments.

Further analysis of our model shows that the spatial correlation for morphing maps with different anchors does not quickly drop to zero (Fig. 3f). This suggests that the spatial correlation may be a feature that bats use to maintain their orientation and place fields across different environments.

Finally, a principal component analysis (PCA) of the population code matrix from morphing maps with different numbers of anchors (2, 5, 15, and 35) reveals that the geometry of bat maps is consistent with the high-dimensional geometry of morphing maps with 35 anchors (Fig. 3g). This suggests that bats might use a high number of anchor points to construct their spatial representations, which allows them to navigate complex environments effectively.

These findings collectively support our hypothesis and suggest a sophisticated neural mechanism by which bats, and potentially other animals, navigate large and gradually changing environments. Future work should aim to experimentally validate the predicted hyperbolic geometry of place field organization and to test whether the number of anchor points used by bats is indeed around 35, as suggested by our PCA analysis.

Adding to our analysis, we also investigated how the optimal number of anchors in our morphing map model is affected by measurement accuracy. This aspect is particularly crucial, considering that real-world measurements of bat place fields are likely to include some degree of noise.

We used the Wasserstein distance metric, a statistical measure used to quantify the distance between two probability distributions. A Wasserstein distance of zero would indicate that the two compared distributions are identical. We computed the Wasserstein distances between the distributions derived from bat data and morphing map simulations for both place field size (Fig. 4a) and the number of place fields per neuron (Fig. 4b).

**Figure 4:**
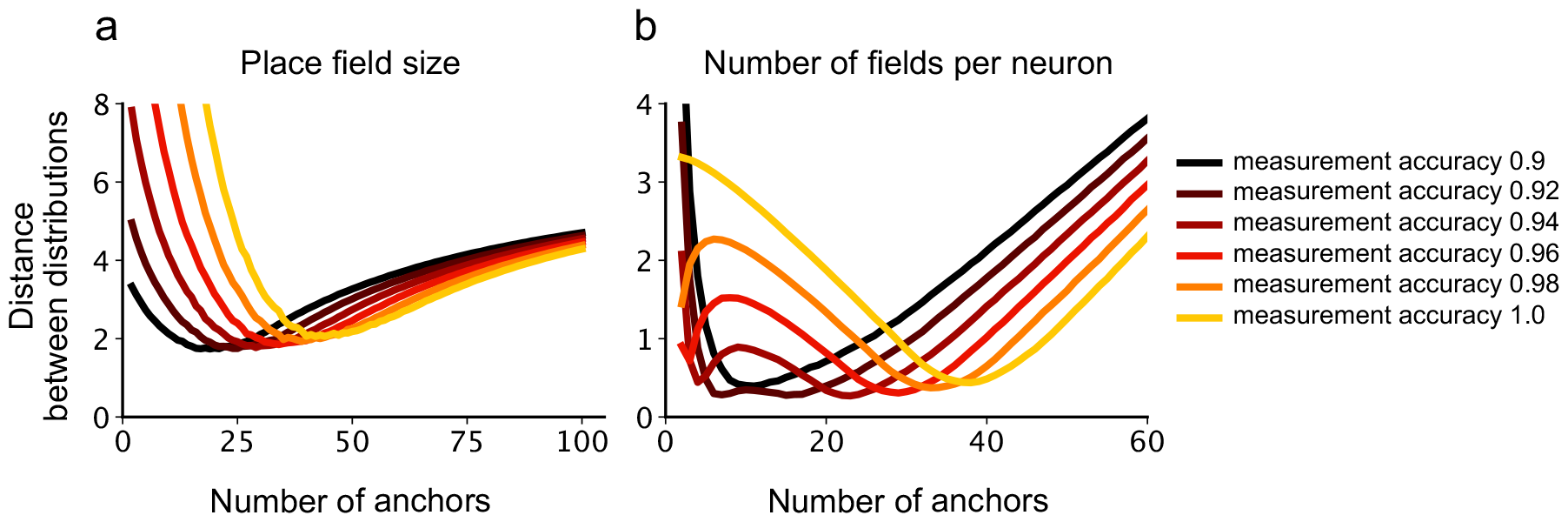
Optimal number of anchors depends on measurement accuracy. Wasserstein distance between distributions derived from bats and morphing map simulations for (a) place field size and (b) the number of place fields per neuron. A distance of zero indicates the two distributions are identical. Analogous to noise that may be introduced during measurements of the bat place fields, the simulated place field maps of 0s and 1s are corrupted by noise, as shown by the different colored curves. For each measurement accuracy there is an optimal number of anchors that gives the best match to the data. As more noise is added, the optimal number of anchors decreases.

To emulate the potential noise that might occur during the real-world measurement of bat place fields, we introduced varying levels of noise into our simulated place field maps of 0s and 1s. The results, illustrated by different colored curves in Fig. 4, reveal an intriguing relationship between measurement noise and the optimal number of anchors.

For each level of measurement accuracy, i.e., each noise level, there exists an optimal number of anchors that minimizes the Wasserstein distance and thus best matches the bat data. As more noise is introduced, the optimal number of anchors decreases, which suggests a trade-off between the complexity of the morphing map model (as reflected by the number of anchors) and the accuracy of the measurements.

This analysis highlights the adaptability of the morphing map model to variations in measurement accuracy. Further experimental studies will be necessary to validate this prediction and to refine our understanding of the interplay between noise, the number of anchors, and the accuracy of spatial navigation in bats.

### Morphing maps generically generate place cell population codes with hyperbolic geometry

It was recently shown [26] that CA1 place cells of the rat hippocampus exhibit non-linear hyperbolic geometry. Through theoretical analysis, Zhang et al. [26] showed that the exponential scale of this representation yields greater positional information than a linear scale.

We see signatures of hyperbolic representation in the morphing map Fig. 3e and in bats Fig. 3e. Thus, we followed [26] and used topological data analysis, specifically the use of clique topology, to quantify the geometry of the codes generated by the morphing map.

Clique topology [27] is topological data analysis method developed to study the geometry of a population of neural recordings *without a priori knowledge of the relevant stimuli being encoded*. It uses pairwise neural correlations and instead of relying on eigenvalues and principal components, an ‘order complex’ is constructed. This is a set of graphs which preserves information about the order of the correlation matrix element magnitudes. Next, cliques (fully-connected subgraphs) of the order complex are examined: the arrangement of cliques which encircle holes (a ‘cycle’) are counted. The Betti curves *β*_1_(*ρ*), *β*_2_(*ρ*) and *β*_3_(*ρ*) count the number of 1-cycles, 2-cycles and 3-cycles respectively. These curves form a topological barcode for the geometry of the neural population [27].

Experimental Betti curves are well matched by sampling from 3D hyperbolic geometry, as confirmed by [26]. To match the Betti curves generated by the morphing map, we sampled points from 3D hyperbolic geometry of different radii and calculated the Betti curves for these samples. To estimate the best matching radius, we used the metric of [26].

To sample points from 3D hyperbolic geometry, we sample points from 3D euclidean space with the probability distribution *p*(*r*) ∝ sinh^2^(*r*). The distance between two points **r**_**1**_ and **r**_**2**_ in hyperbolic space is given by

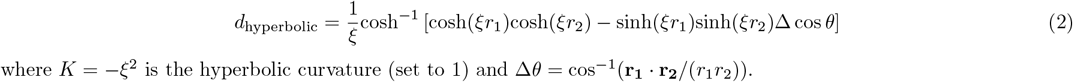

where *K* = −*ξ*^2^ is the hyperbolic curvature (set to 1) and Δ*θ* = cos^*−*1^(**r**_**1**_ · **r**_**2**_*/*(*r*_1_*r*_2_)).

### Number of anchor points controls hyperbolic radius

The hyperbolic geometry we observe is a function of the number of anchor points used in the morphing map. The number of anchor points effectively controls the complexity of the resulting hyperbolic geometry. As seen in Fig. 5c, as the number of anchors increases, the optimal hyperbolic radius also scales. This highlights the versatility and adaptability of the morphing map in response to increasing the complexity of the input data.

**Figure 5:**
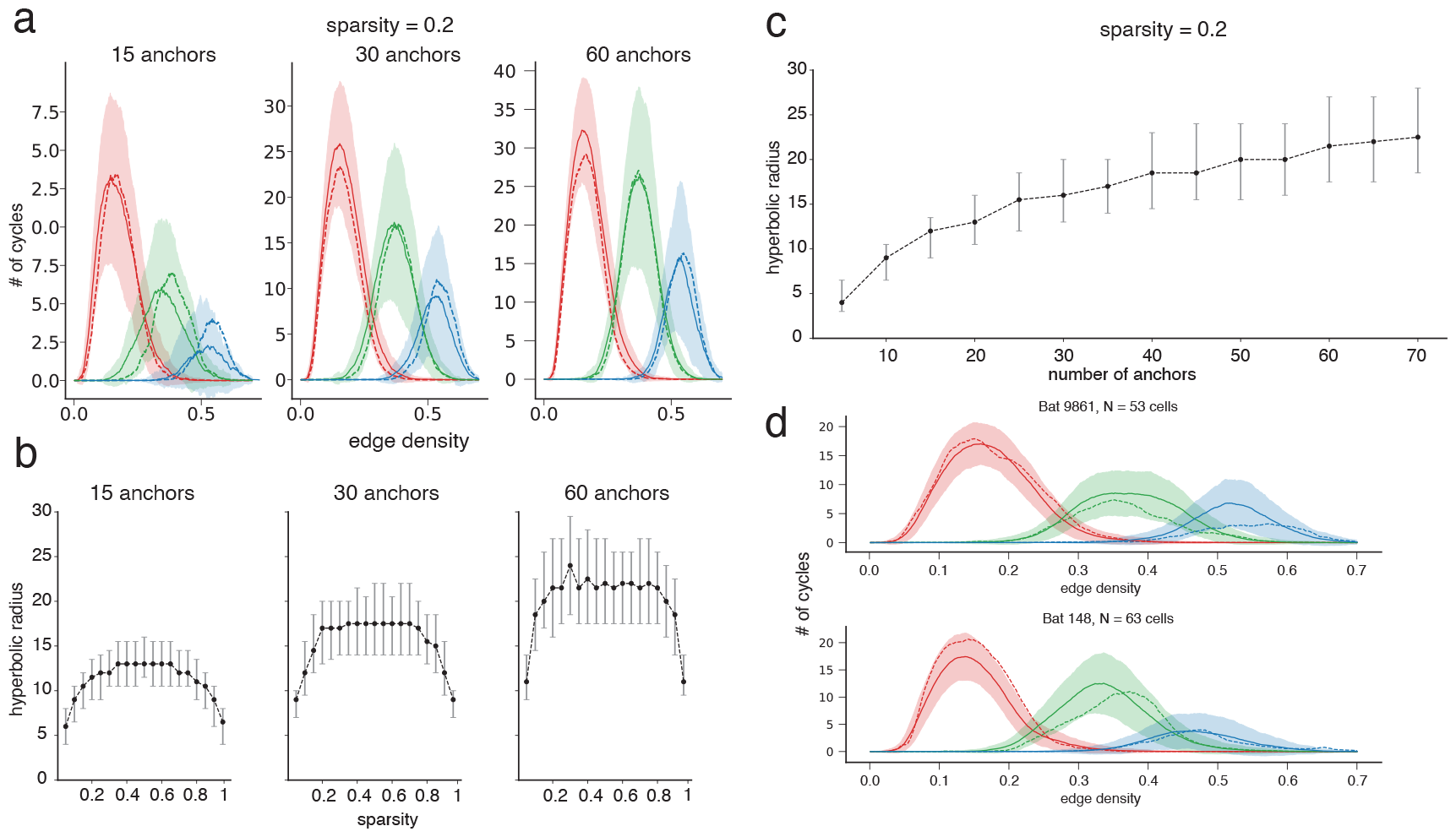
Morphing maps create population codes that exhibit hyperbolic geometry whose radius scales with number of anchors. (**a**) The Betti curves *β*_1_(*ρ*), *β*_1_(*ρ*) and *β*_3_(*ρ*) for a morphing map with 15, 30, and 60 anchor points respectively. The dashed line denotes the Betti curves of hyperbolic space with the best matching radius (shading indicates mean ± s.d.). (**b**) For a given number of anchors (15, 30 and 60 shown), hyperbolic radius varies with the sparsity of the morphing map, peaking at a sparsity of 50%. (**c**) The optimal hyperbolic radius scales with the number of anchors. (**d**) Average Betti curves *β*_1_(*ρ*), *β*_2_(*ρ*) and *β*_3_(*ρ*) for 2 bats along with those generated with morphing map with 35 anchor points.

In addition, we studied the effect of sparsity in the morphing map on the hyperbolic geometry. The sparsity of the morphing map significantly influenced the hyperbolic radius as shown in Fig. 5b. We observe that as the map becomes denser (less sparse), the hyperbolic radius increases. This reflects the intuitive expectation that more densely connected neural populations can support a richer geometric structure.

### Comparison with bats

The comparison between the hyperbolic geometry observed in bats and that generated by the morphing maps revealed a striking similarity. The average Betti curves obtained from both bats’ neural data and the morphing maps exhibited a significant overlap, suggesting a shared underlying hyperbolic geometric structure. To achieve this averaged representation, the process involved randomly selecting 80% of the available data and repeating it 300 times. Subsequently, the Betti curves were computed for each iteration. For a visual comparison, please refer to Figure 5d.

This compelling resemblance between the results derived from the morphing maps and the neural data of bats lends support to the hypothesis that morphing maps can generate neural population codes with inherent hyperbolic geometry, potentially reflecting how biological brains represent spatial information. However, additional experimental and theoretical investigations are necessary to establish this connection conclusively.

### Investigating position decoding and associated errors in the morphing map model and the bats

In this section, we aim to simulate and better understand the complex navigational strategies of bats by examining models with various numbers of anchor points and adjusting the probability of neuronal firing.

To mimic the patterns of neuronal activity, we begin with a given place field map, *M*_(*l,n*)_, and generate a raster plot, *r*_(*l,n*)_. Here, *l* is the location and *n* is the neuron. Neurons with a place field at a certain location fire with a probability *p*_*firing*_, while the remainder remain inactive.

We then proceed with position reconstruction and error calculation within our model. We define the global error, Δ, as the absolute difference between the real location, *x*_*l*_, and the reconstructed location, 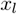, averaged across all locations:

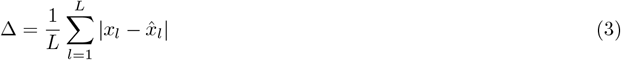

To determine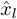, we compute the likelihood of observing a specific firing pattern at each location, and then take the location that maximizes the likelihood.

The probability of observing a pattern of 0s and 1s across *N* neurons at a specific location *l* is determined based on prior knowledge about whether or not each neuron has a place field at this location. This information is stored in the binary map *M*_(*l,n*)_, where *M*_(*l,n*)_ = 1 if neuron *n* at location *l* has a place field, and *M*_(*l,n*)_ = 0 if it does not. If a neuron has a place field then it has a probability *p*_*firing*_ of firing, i.e., outputting a 1. If a neuron has no place field then it cannot fire.

The firing pattern of neurons is modeled as a Bernoulli process, where each neuron *n* at location *l* either fires (outputting a 1) or does not fire (outputting a 0). Thus, the probability *P* (*n* = *r*|*l*) of that neuron *n* firing or not *r* ∈ *{*0, 1*}* at location *l* can be expressed as

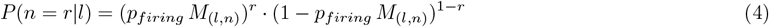

Given the empirically observed activity of N neurons 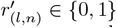 where *n* = 1, 2, …, *N*, and assuming independence among neurons, we can compute the log-likelihood of this data at each location *l* as

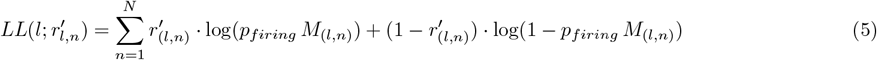

This log-likelihood measure quantifies the fit between the observed firing patterns and those predicted by the model, under the stated assumptions. The second term in equation 5 accounts for neurons that did not fire when there was a corresponding place field.

For each firing pattern, 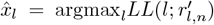 is the location that maximizes the log-likelihood. This approach provides a robust method for position reconstruction based on patterns of neuronal activity, aiding in our understanding of bat navigational strategies.

Our model enables us to investigate the performance of different morphing maps, each characterized by a unique number of anchor points. As demonstrated in Fig. 6, as the probability of neuronal firing increases, a morphing map with a larger number of anchor points (35 in our experiments) consistently generates non-local reconstruction errors, implying a distributed spatial representation. Conversely, a map with fewer anchor points (2 in our case) produces local errors, suggesting a more localized spatial representation. These findings echo the spatial representation in bats’ navigation, further affirming the high-dimensional geometric consistency between our model and bats’ real-world navigation strategies.

**Figure 6:**
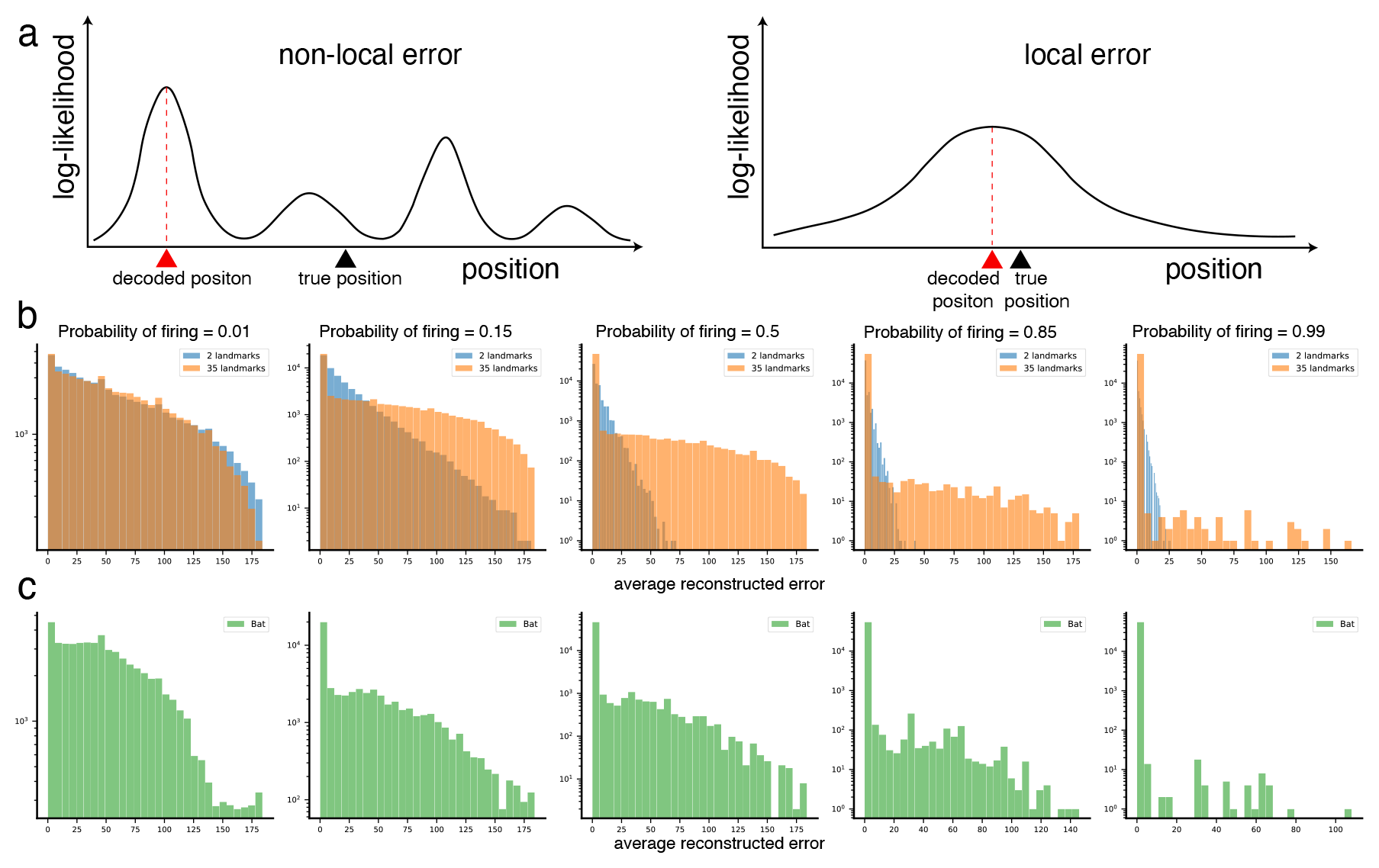
Errors made in decoding position from the morphing map place cell population code. (**a**) Schematic of non-local and local errors. (**b**) Average reconstruction error from 300 morphing maps, with 2 (blue) and 35 (orange) anchors each. (left to right) As the probability of firing increases, the errors made by the 35 anchor map remain non-local while the errors made by the 2 anchor map are local. **(c)** Analogous to above, but for the place cell population code from bat 9861.

Moreover, we analyzed the influence of measurement accuracy on the reconstruction error. The results, illustrated in Fig. 7, are averages from 300 different simulations of morphing maps, each containing 35 anchor points and a sparsity level that mirrors the bats’ neuronal activity sparsity. The scenarios spanned a range of firing probabilities — from a relatively low *p*_*firing*_ = 0.15 to an elevated *p*_*firing*_ = 0.85. Our analyses indicated that the decoding model performance remains consistent across variations in measurement accuracy, regardless of the firing probability. This robustness against potential inaccuracies in measurements underscores the model’s reliability. The minimal influence of measurement accuracy on the decoding error further highlights the effectiveness of our model in accurately decoding the complex patterns of neuronal activity and spatial representations observed in bats, even when subjected to multiple simulations with varying sparsity and anchor points.

**Figure 7:**
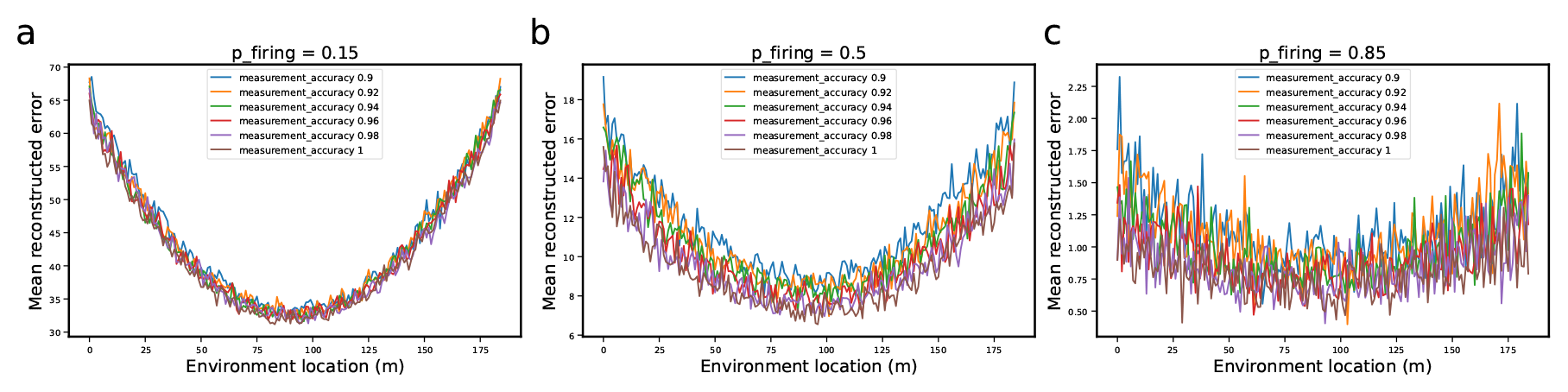
Evaluation of measurement accuracy’s impact on error reconstruction across diverse firing probabilities, based on multiple simulations. Each curve represents the average reconstructed error per location from 300 different generated morphing maps, each containing 35 anchor points and a sparsity level matching that of the bats. Three firing probability scenarios were investigated: (**a**) *p*_*firing*_ = 0.15. (**b**) *p*_*firing*_ = 0.5 (**c**) *p*_*firing*_ = 0.85. Irrespective of the firing probability, changes in measurement accuracy did not significantly influence the error reconstruction, indicating a robust decoding performance across different degrees of measurement accuracy.

### Predictions

#### Neural population level metrics

We use the Procrustes distance [28, 29], a measure of shape difference, to evaluate the similarities and differences between these maps, considering several factors such as the number and location of anchor points.

Two bats, flying through the same tunnel, present an interesting scenario for analysis. Their place field map distances are illustrated in blue in Fig. 8. To interpret these results, we also compare the distances with those generated by our model. We simulate two distinct situations, shown by gray and red curves in the figure.

**Figure 8:**
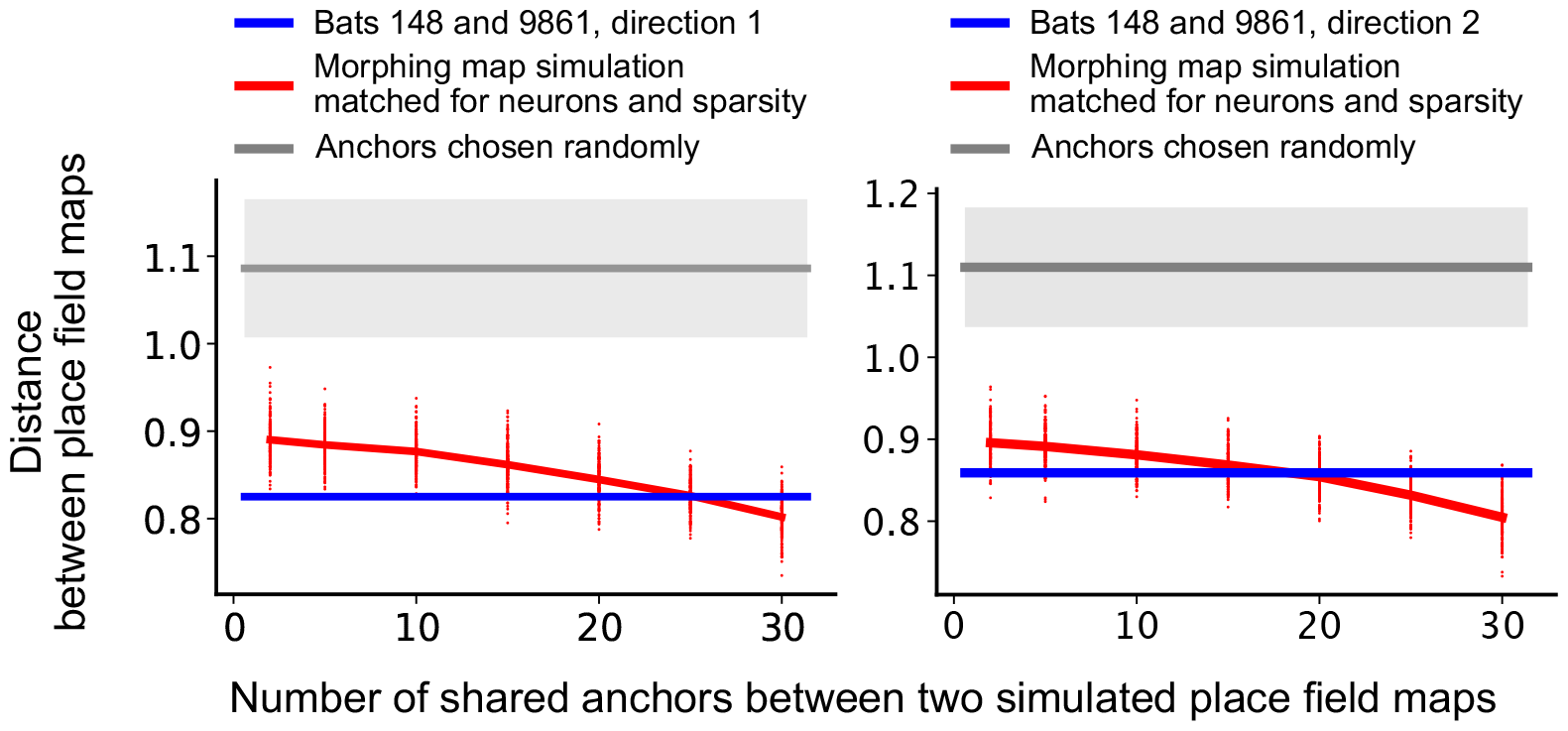
Predicting shared structure between bat place field maps. Procrustes distance between place field maps from two bats flying through the tunnel in direction 1 (left) and direction 2 (right) are shown in blue. The gray curve shows the distance between two simulated place field maps when the number and location of anchor points are chosen randomly for each map (error bars show one standard deviation). This contrasts with the red curve where both simulated place field maps have a fixed number of anchor points (30 in this example). Each red dot in the figure is the distance between two simulated place field maps having 30 anchor points, but the number of shared anchor points is varied (x-axis). The location of the anchor points are also varied, leading to a spread in distances. The simulated place field maps for both the gray and red curves have the same number of neurons and sparsity as the bat data, and we include anchor points at both ends of the environment. The distance between the bat place field maps (blue) is less than the distance between random place field maps (gray) suggesting there may be shared structure between the place field maps for the two bats. Place field maps that look random at the level of an individual bat may actually have shared structure when comparing multiple bats. More specifically, if two bats share some anchor points then, according to the morphing map simulations, the place field maps have a lower Procrustes distance than if the anchor points for both simulated bats are chosen independently. This is shown by the red curve; as the number of shared anchor points increases the distance decreases. Using the morphing map simulations we can estimate the number of shared anchor points for the two bats by noting the intersection of the red and blue curves. This illustrates one way in which the morphing map simulations can be used to make predictions, however, these may not be feasible in practice as accurate predictions depend on the measurement error and the number of anchor points, which may be unknown.

In the first scenario, represented by the gray curve, we randomly choose the number and location of anchor points for each simulated map. This produces a wider range of distances (error bars indicate one standard deviation).

In contrast, the second scenario, depicted by the red curve, uses a fixed number of anchor points (30, in our example) for both maps. Each point on the red curve indicates the distance between two simulated maps with 30 anchor points, as we alter the number of shared anchor points and their locations.

Interestingly, the distance between the bat place field maps (blue curve) is smaller than the distance between randomly generated place field maps (gray curve). This suggests that despite the maps looking random at the level of an individual bat, there might be some shared structure when comparing maps from multiple bats.

According to our model, if two bats share some anchor points, their place field maps tend to have a lower Procrustes distance compared to the case when the anchor points for both simulated bats are chosen independently. This is demonstrated by the decreasing trend in the red curve as the number of shared anchor points increases.

Our morphing map simulations predict that the number of shared anchor points between two bats could be inferred from the intersection of the red and blue curves. This offers a glimpse of how the morphing map model can be used to make predictions about complex biological phenomena. However, these predictions should be interpreted with caution, as they are sensitive to factors such as measurement error and the number of anchor points, which may not always be known. This section, therefore, underlines the potential of the morphing map model as a predictive tool, even though certain constraints.

#### Anchor point ablation and displacement

Our exploration extended to examining the impact of anchor point removal from neural maps, as shown in Fig. 9. We first generated an original neural map with 166 neurons and 5 anchor points. This original map was then regenerated a thousand times; each time it was regenerated all the anchor points were the same but the place fields at intermediate locations were filled in stochastically according to the morphing map algorithm. The regenerated map was then compared to the original map and the absolute difference in the number of place fields was computed to create a control curve as shown in blue in Fig. 9a. The original map and the regenerated maps both share the same anchor points so the difference in the number of place fields is zero at the anchor points, i.e. the control curve, shown in blue, is zero at these locations. Vertical dashed lines indicate the locations of anchor points.

**Figure 9:**
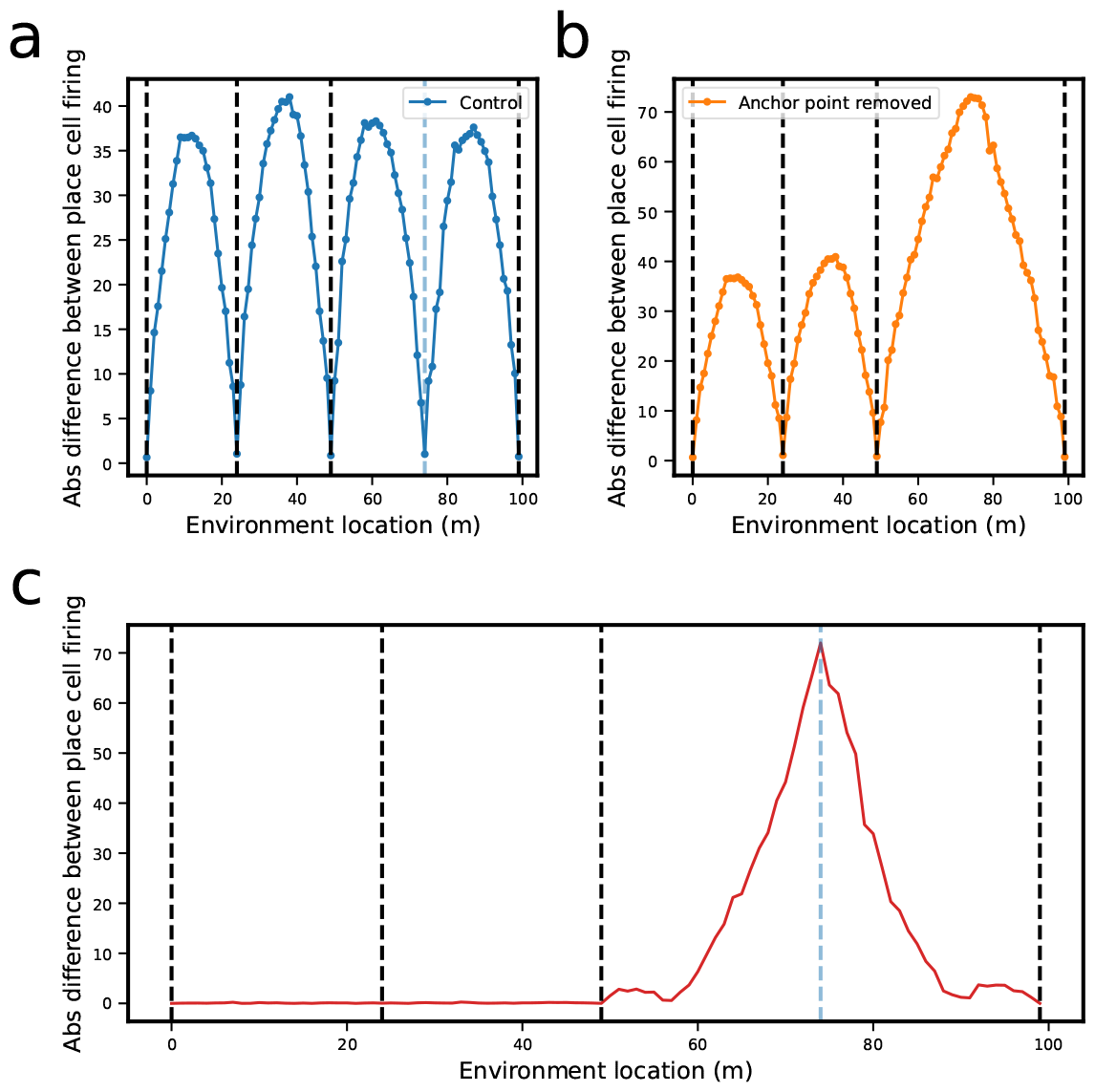
Effects of anchor point removal on morphing maps. An original morphing map was created with 166 neurons and only 5 anchor points for simplicity. To establish a baseline for comparison, the original map was then regenerated a thousand times while maintaining the neural firing rates at anchor points to form a set of control maps. Concurrently, an experiment was conducted where one anchor point was randomly removed. This altered map was regenerated a thousand times, each time excluding the removed anchor point. The figure compares three map scenarios - the original map, the control maps (containing all the same anchor points as the original), and the maps with the removed anchor point. Dashed vertical lines indicate the locations of anchor points. (**a**) The absolute difference in the number of place fields between the original map and the regenerated control maps (blue). (**b**) Comparison between the original map and regenerated maps that have one anchor point removed (orange). (**c**) The absolute difference between the control maps and the maps with the removed anchor point. Notably, the removal of an anchor point induces characteristic differences at the location of the removed anchor point in the regenerated map. The bell shape is caused by the stochasticity of turning off or on a place field in the path between landmarks.

Simultaneously, we emulated a scenario where a single anchor point was randomly selected and removed. This modified map was also regenerated a thousand times, each iteration excluding the selected anchor point. In Fig. 9b we generated morphing maps after removing a single anchor point. This modified map was regenerated a thousand times, each iteration excluding the same anchor point. Each regenerated map was then compared to the original map and the absolute difference in place cell firing rates between maps was computed to create the curve shown in orange (panel b). Fig. 9c shows the absolute difference between the control maps used in panel (a) and the maps missing an anchor point. The bell shape is caused by the stochasticity of turning off/on a place field in the path between landmarks.

Interestingly, the removal of an anchor point results in unique differences at the location of the removed anchor point in the regenerated map. This observation provides significant insight into the role of anchor points in maintaining the structure and stability of neural maps.

Fig. 10 shows a similar set of analyses except we now generated morphing maps after shifting an anchor point half the distance closer to one end. The results of Fig. 9 and Fig. 10 serve as experimentally testable predictions of the morphing map model.

**Figure 10:**
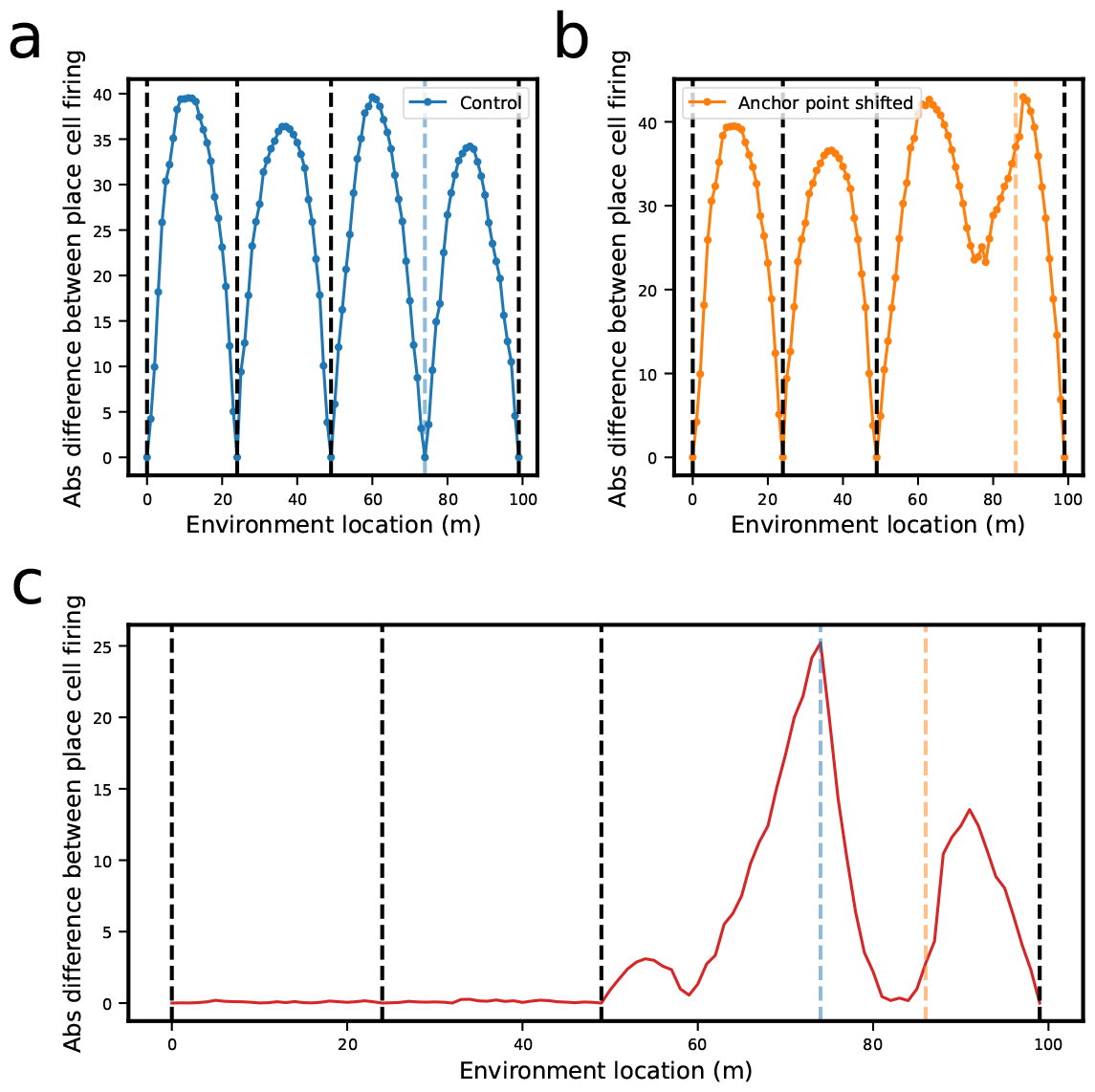
Effects of a shifting anchor point on morphing maps. An original morphing map was created with 166 neurons and only 5 anchor points for simplicity. To establish a baseline for comparison, the original map was then regenerated a thousand times while maintaining the neural firing rates at anchor points to form a set of control maps. Concurrently, an experiment was conducted where one anchor point was shifted half the distance closer to one end. This altered map was regenerated a thousand times, each time with the shifted anchor point. The figure compares three map scenarios - the original map, the control maps (containing all the same anchor points as the original), and the maps with the shifted anchor point. Dashed vertical lines indicate the locations of anchor points. (**a**) The absolute difference in the number of place fields between the original map and the regenerated control maps (blue). (**b**) Comparison between the original map and regenerated maps that have one anchor point shifted (orange). (**c**) The absolute difference between the control maps and the maps with the shifted anchor point.

## Discussion

In this study, we have introduced a novel morphing map algorithm inspired by the spatial navigation strategies of bats, which incorporates hyperbolic geometry to model place fields within the hippocampus. Our findings reveal a complex, scalable spatial representation that suggests place cells adapt to large environments by integrating landmark-based navigation with inherent geometric principles. This approach contrasts with previous models focused on Euclidean space, offering a more nuanced understanding of spatial memory and navigation in extensive areas.

The implications of our research extend beyond the biological significance, hinting at potential applications in developing more sophisticated navigational algorithms for robotics and AI, which could benefit from incorporating principles derived from natural navigation strategies. Moreover, our work prompts a reevaluation of how spatial information is encoded across different species, supporting the notion that spatial cognition may involve a blend of geometric frameworks tailored to the environmental context.

However, the study’s limitations, including the reliance on bat hippocampal data and the specificity of the modeled environment, suggest the need for further research. Future studies could explore the applicability of the morphing map algorithm across a broader range of species and environmental conditions, potentially offering insights into the evolution of navigational strategies and the neural basis of spatial memory disorders.

In conclusion, our research contributes a pivotal step towards understanding the intricate mechanisms underlying spatial representation in large environments, bridging the gap between biological insights and technological applications. This study not only enhances our comprehension of hippocampal function but also opens new avenues for interdisciplinary research, combining neuroscience, mathematics, and computer science to unravel the complexities of cognitive mapping.

## Acknowledgments

We thank Tamir Eliav and Nachum Ulanovsky for generously sharing the experimental data, and Tatyana Sharpee and Huanqiu Zhang for providing the Matlab code necessary to produce the Betti Curves. We also extend our thanks to Nachum Ulanovsky, Misha Tsodyks, Alessandro Treves, and Mikail Katkov for useful discussions. MN, MK, CC thank the K. Lisa Yang ICoN center at the McGovern Institute, MIT for funding support.

## Notes

### Competing Interest Statement

The authors have declared no competing interest.

